# Attitudes Toward Cougar Restoration in Seven Eastern States

**DOI:** 10.1101/2023.05.26.542322

**Authors:** Shelby C. Carlson, John A. Vucetich, Lexi Galiardi, Jeremy T. Bruskotter

## Abstract

**BACKGROUND:** Cougars (*Puma concolor*), also known as mountain lions, pumas, catamounts, and Florida panthers, once ranged widely throughout the United States (McCollough, 2011). Following intensive human persecution, anthropogenic-driven habitat loss, and unrestricted hunting of the prey species upon which cougars depend, populations were extirpated from much of the country (Cardoza & Langolis, 2002). In the northeastern U.S. specifically, cougars were functionally extinct by the early 1900s. Recent research assessing potential habitat for cougars suggests numerous areas exist to restore the species throughout portions of their historic eastern range (Winkel et al., 2022; Yovovich et al., 2023). But are the humans inhabiting this region today supportive of cougar restoration?

**STUDY OBJECTIVES:** The primary goal of this study was to make a preliminary assessment of support for cougar restoration at the state level in several states deemed to have substantial habitat for cougars (i.e., Maine, Massachusetts, New Hampshire, New York, Pennsylvania, Vermont, and West Virginia). Additionally, we sought to identify individual-level correlates of support for, and opposition to, cougar restoration.

**STUDY FINDINGS:** Results from an online survey of residents of seven eastern states with potential cougar habitat (n=2756) suggest that support for cougar restoration is much higher than opposition to cougar restoration. Ratios of strong support to strong opposition range from approximately 4:1 to 13:1. Maine has the highest ratio of strong support to strong opposition at 13:1, indicating that for every one person opposing cougar restoration in the state, we can expect 13 people to support it. Vermont and New Hampshire have the second highest ratio of strong support to strong opposition at 12:1 each. New York and Massachusetts have the second lowest ratio of strong support to strong opposition, at 5:1 each. West Virginia and Pennsylvania have the lowest ratio of strong support to strong opposition with ratios of 4:1, indicating that for every one person opposing cougar restoration in these states, we can expect 4 people to support it. Results also reveal that states with the lowest ratio of strong support to strong opposition tend to have the highest proportion of respondents expressing neutrality toward the idea of restoration.

At the individual-level, support for cougar restoration was higher among men, respondents identifying “strongly” or “very strongly” as a hunter or a conservationist, those with mutualist wildlife value orientations, urban residents, and respondents identifying as politically liberal.

**IMPLICATIONS:** Given the current structure of wildlife management in the U.S., efforts to restore cougars throughout significant portions of their historic range will depend in large part on actions taken by state wildlife management agencies. Finding support for cougar restoration among many of the constituents for whom state wildlife agencies are expected to operate on behalf of – including both hunters and conservationists – this study offers valuable insights regarding the *social* feasibility of restoring cougars to the eastern U.S.

Importantly, while a majority of respondents were supportive of cougar restoration, a considerable portion of the population in each state expressed neutrality toward the idea of cougar restoration. Extant research from the behavioral sciences suggests these individuals may be more likely to change their attitudes toward cougar restoration in response to new information. Whether any such change would result in greater support or greater opposition toward cougar restoration is likely dependent on several factors, including the way in which information regarding the potential risks and benefits of the species is presented (Slagle et al., 2013), as well as the source/messenger from which new information is provided (e.g., Fielding et al., 2020).

**COVER IMAGE:** Word cloud produced from survey participant responses to free association when they think of cougars.

## METHODOLOGY AND DESIGN

### SAMPLING

We commissioned a survey of adult residents of seven eastern states (Maine, Massachusetts, New Hampshire, New York, Pennsylvania, Vermont, West Virginia) in February 2022. Respondents were recruited through an online panel maintained by the commercial sampling firm, Qualtrics (Provo, Utah) and the survey was delivered via Qualtrics’ online software. Quota-based sampling methods were used for participant selection in our study, with the goal of (i) reaching approximately 400 individuals in each state (with the exception of Vermont) and (ii) obtaining a 1:1 male/female ratio among respondents across the nine states (but not necessarily within each state). The survey was reviewed by The Ohio State University’s Office of Responsible Research Practices and determined to be exempt from Institutional Review (protocol number 2021E1229).

### WEIGHTING

Post hoc weights were created to enhance state-level representativeness of our sample among selected social and demographic characteristics (i.e., sex, educational attainment, and political ideology). State population distributions for sex and educational attainment were based on benchmarks from the 2016-2020 American Community Survey (ACS), conducted by United States Census Bureau (www.census.gov/acs/, date last accessed 10 September 2022). State population distributions for political ideology were based on estimates from the 2018 Gallup U.S. Poll (Jones 2019). Weights were developed using iterative proportional fitting (i.e., raking) and applied via the RAKE extension in IBM SPSS Statistics v28 (Chicago, Illinois).

## STUDY RESULTS

### 1. What actions should state wildlife agencies prioritize?

Participants were presented with a list of actions that might be seen as being within the purview of state wildlife agencies and asked to prioritize that list according to which actions were most important to the respondent. The complete list of actions included:

- Restoration of species that are locally extinct or imperiled
- Increasing opportunities to hunt and/or trap species
- Purchasing or leasing lands to create recreational access
- Management of existing lands to improve habitat
- Removal of invasive or exotic species

Figure 1 (next page) shows the proportion of respondents indicating which activity should be top-ranked. The graph further indicates how participants from the seven eastern states responded to this survey item (n=2756) as well as respondents who also identify “strongly” (n=142) or “very strongly” (n=158) as being hunters.

**Figure 1.**
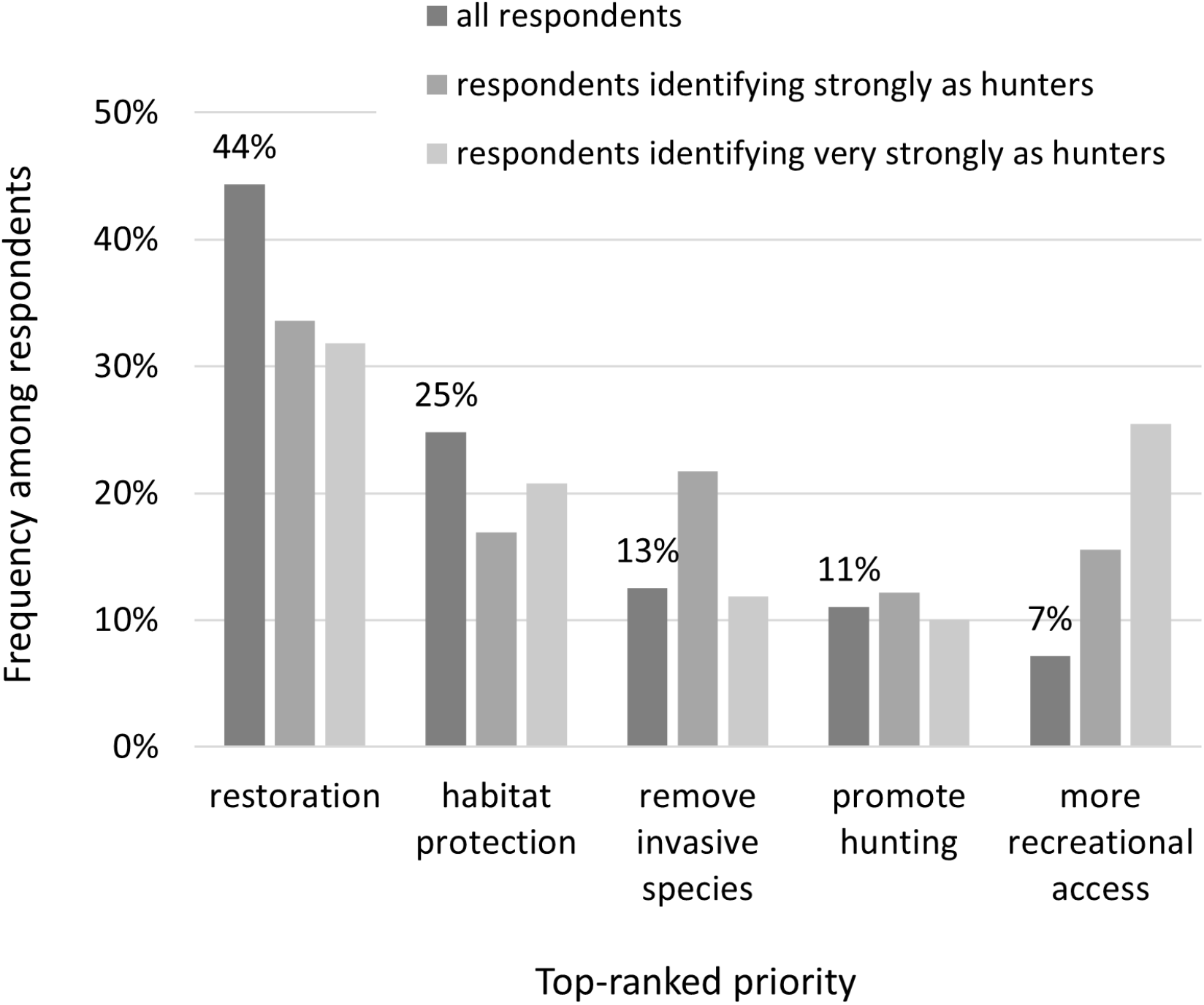
Frequency of respondents giving top-ranking to each of several actions they believe should be prioritized by state fish and wildlife agencies.

**Restoration is the top-ranked priority for residents overall and for those who identified strongly (or very strongly) as hunters**. These results are relevant for: (i) assessing the extent to which the priorities of state wildlife agencies align with the priorities of their constituents, and (ii) identifying activities that are top-ranked by both the hunting community and agencies’ broader constituency.

### 2. Overall attitudes toward cougar restoration

Participants were presented with some basic information about cougars via an infographic (Figure 2). The information described cougars’ geographic range, basic biology, and behavior.

**Figure 2.**
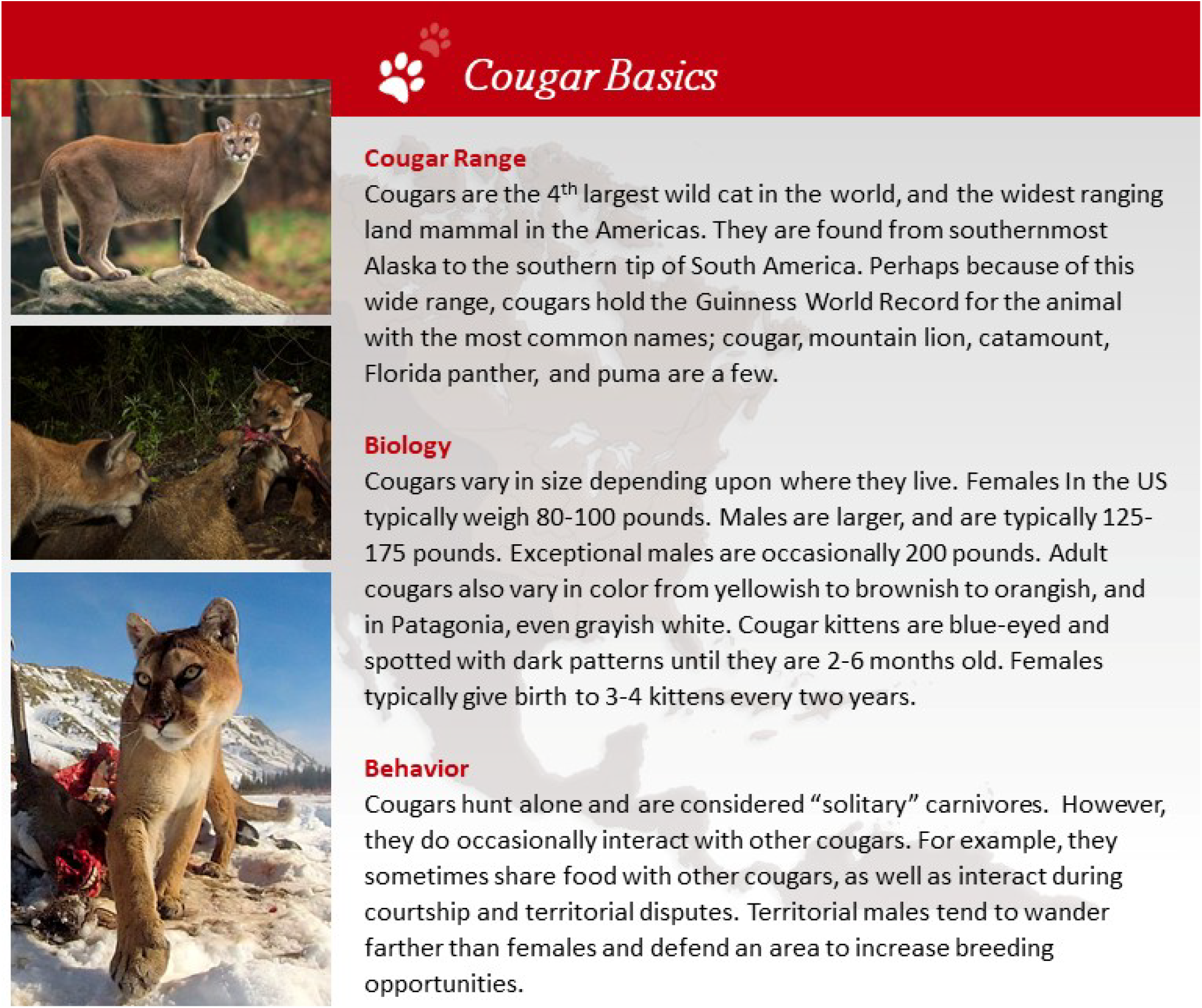
Participants were shown general information about cougars, including their range, biology, and behavior in the form of this (above) infographic.

Following the presentation of that information, participants were asked to record, in a few words, the first thought or image that comes to their minds when they think about cougars. The results of that query are informally represented by the word cloud that is the cover page of this report.

Participants were also asked to “indicate whether you would support or oppose reintroduction of” cougars and were presented with this survey item:

If cougars were reintroduced to rural areas of the eastern United States, what would be your reaction?

- This is a GREAT idea and I would express my support
- This is a GOOD idea and I don’t care enough to get involved
- I don’t have an opinion / I am not sufficiently informed to judge
- This is a BAD idea, but I don’t care enough to get involved
- This is a TERRIBLE idea and I would express my opposition

The distribution of responses (n=2756) to those two survey items for the seven eastern states (MA, ME, NH, NY, PA, VT, WV) is shown to the right in Figure 3.

**Figure 3.**
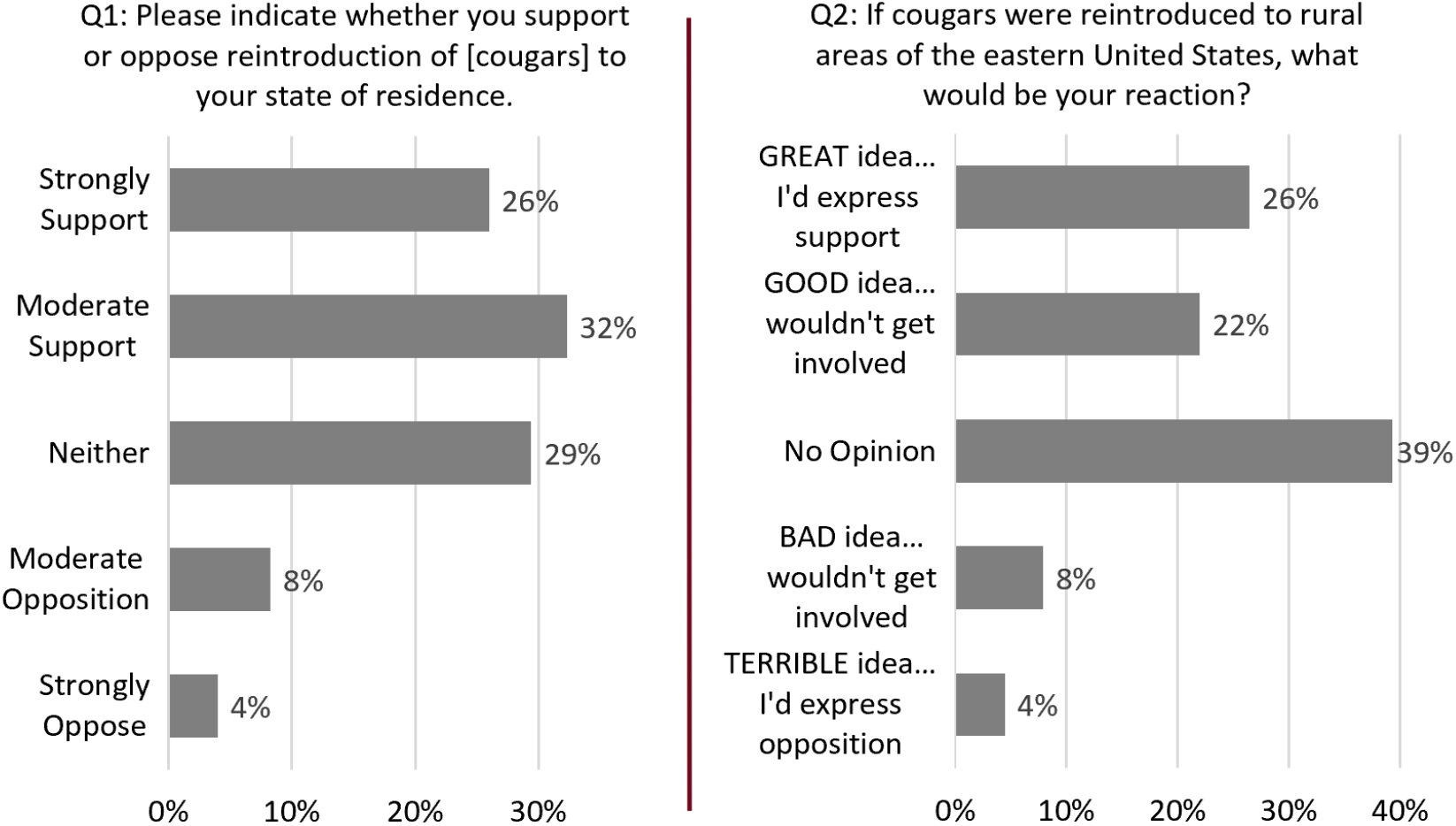
Attitudes toward cougar restoration as indicated by two survey items.

In analyzing survey data, it is common practice to create aggregate measures by averaging responses to similarly worded items that are strongly correlated. Doing so mitigates measurement error associated with differences in finer details of the wordings used in each survey item (DeVellis & Thorpe, 2021). To calculate an average, we converted the text responses to numerical values (-2, -1, 0, 1, 2), where -2 represents “strong opposition” or “Terrible idea.” After averaging the responses, we coded respondents according to the table below. The results are shown in the graph below (Figure 4).

**Figure 4.**
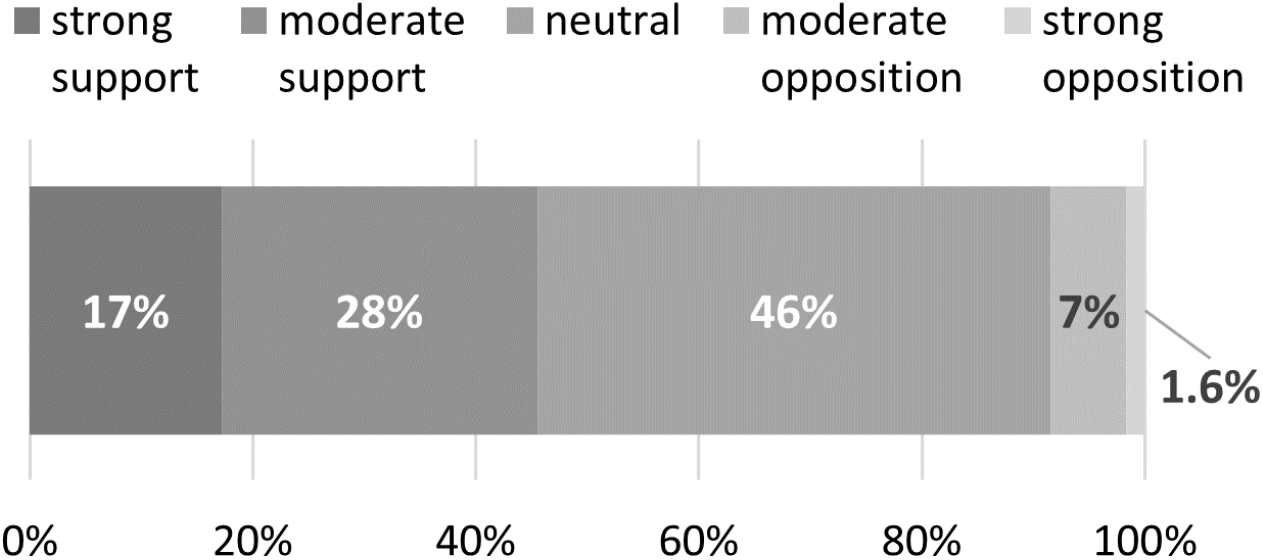
Distribution of attitudes for cougar restoration as indicated by a combined (averaged) measure.

| Score | Classification |
| --- | --- |
| -2 | Strong opposition |
| -1.5, -1.0 | Moderate opposition |
| -0.5, 0, +0.5 | Neutral |
| 1.0, 1.5 | Moderate support |
| 2 | Strong support |

While the largest share of people fall in the ‘neutral’ category, the ratio of strong support to strong opposition is approximately 10:1, meaning that for every person who strongly opposes cougar restoration one can expect approximately 10 people who strongly support their restoration. Results presented in sections 3 through 9 are based on this restoration scale. Results presented in section 10 are based on Q2 (where the response set includes key words, GREAT, GOOD, etc.).

### 3. Attitudes about cougar restoration among hunters and non-hunters

Toward the end of the survey, participants were asked to indicate the degree to which they identify as a hunter (not at all, slightly, moderately, strongly, very strongly). The graph below (Figure 5A) shows the association between intensity of one’s identity as a hunter and attitudes toward cougar restoration.

**Figure 5.**
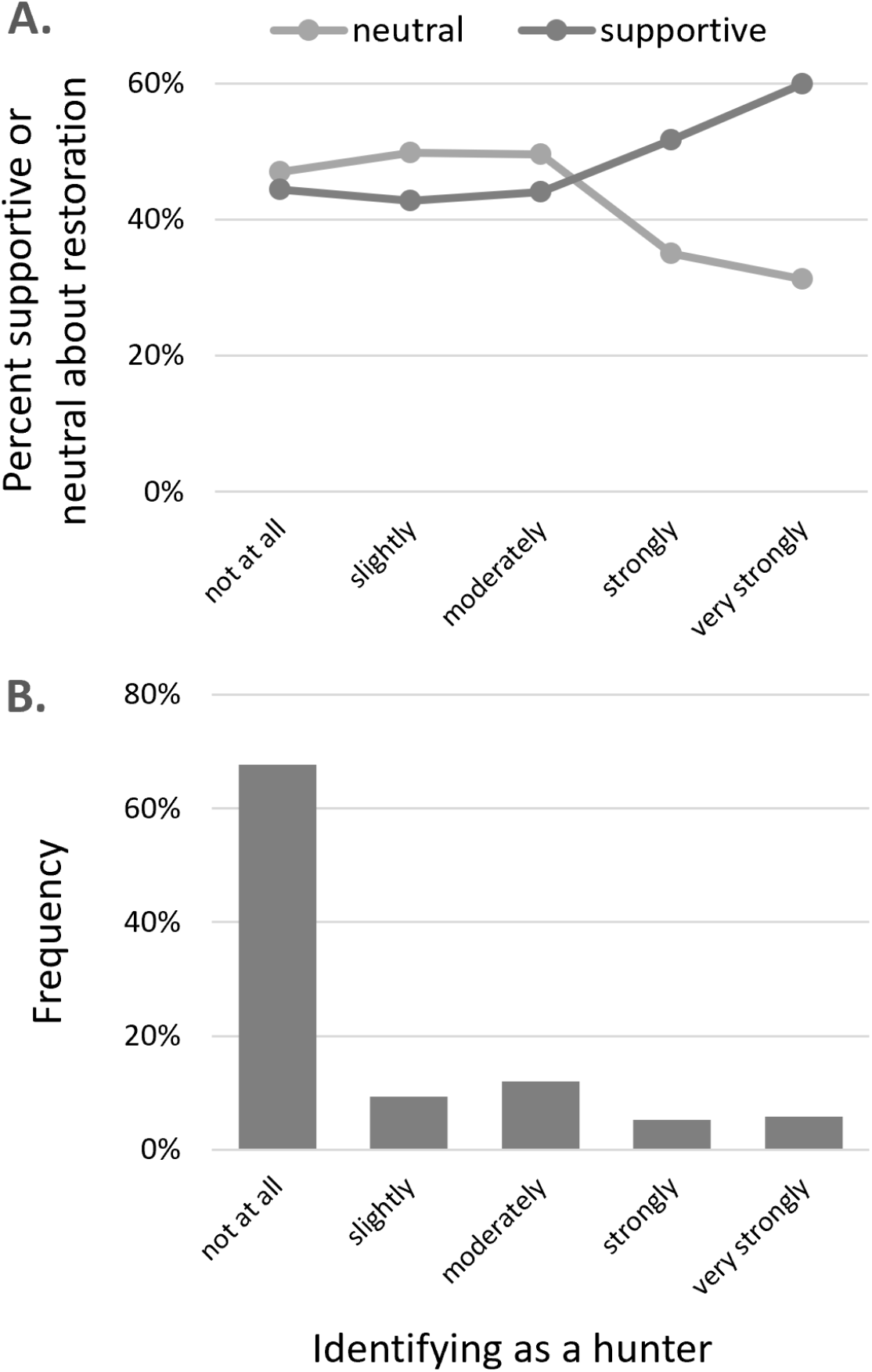
Attitudes about cougar restoration among those identifying to varying degrees as a hunter.

The lower panel (Figure 5b) of this graph provides additional context by indicating the portion of the sample (for the seven eastern states) who identify to varying degrees as being a hunter.

Attitudes about cougar restoration are similar among three groups of people, i.e., those who identify with hunting “not at all,” “slightly,” or “moderately.”

Among those who identify “strongly” or “very strongly” as hunters the frequency of support for cougar restoration is higher and the frequency of being neutral toward cougar restoration is much lower.

For context, these differences in support and neutrality represent a relatively small portion of the entire sample – as only about 10% of respondents identify strongly or very strongly as hunters. The decline in neutrality is also associated with a large increase in strong support and a small increase in strong opposition to cougar restoration.

### 4. Attitudes about cougar restoration among self-identified conservationists and non-conservationists

Survey participants were also asked to indicate the degree to which they identify as a conservationist (not at all, slightly, moderately, strongly, very strongly). The graph below (Figure 6A) shows the association between intensity of one’s identifying as a conservationist and attitudes toward cougar restoration.

**Figure 6.**
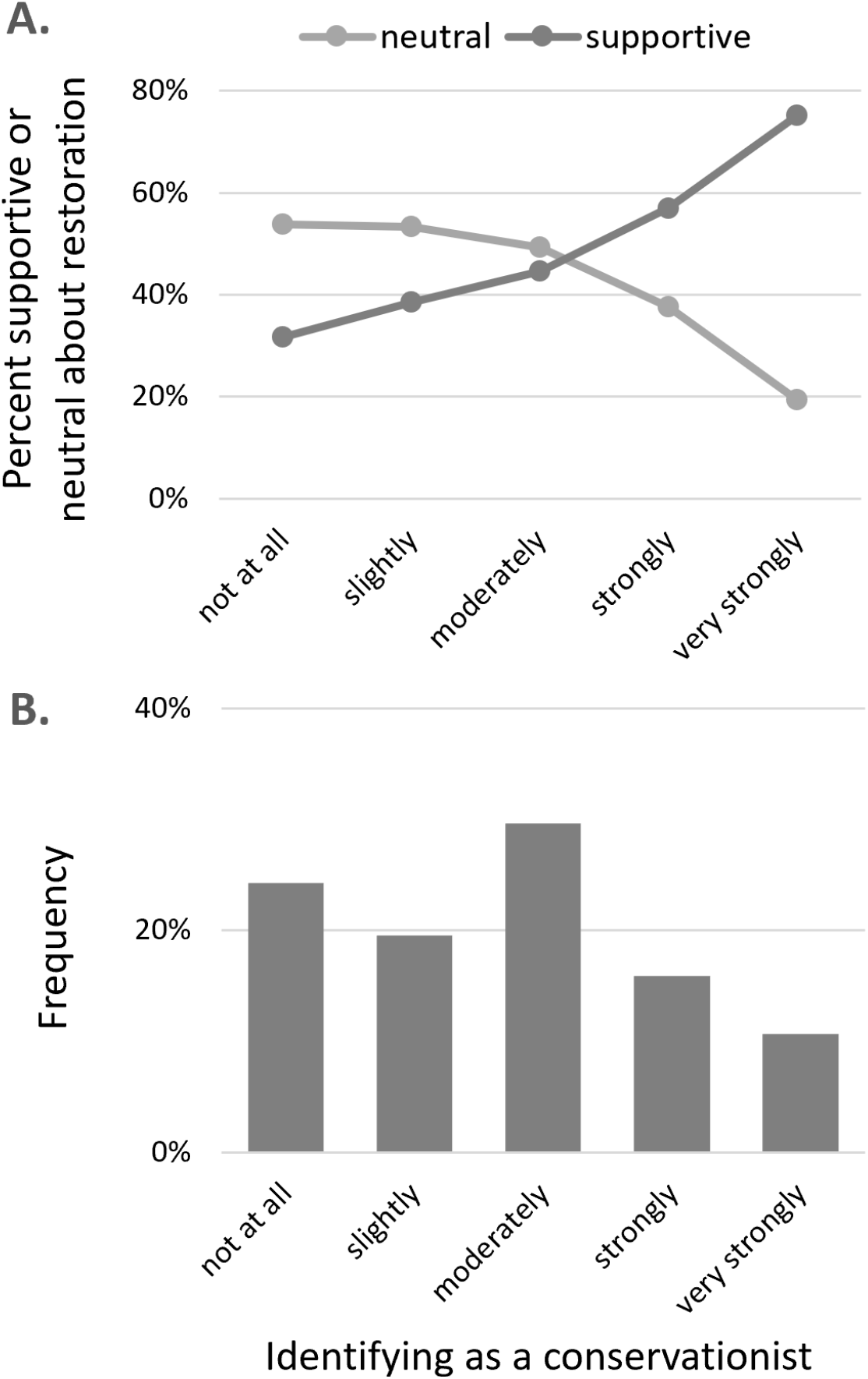
Attitudes about cougar restoration among those identifying to varying degrees as a conservationist.

Attitudes about cougar restoration become increasingly supportive as one’s identity is increasingly associated with being a conservationist. Concomitantly, the frequency of neutral views toward cougar restoration declines among those who more strongly identify as being a conservationist.

The lower panel (Figure 6B) of this graph provides additional context by indicating the portion of the sample (for the seven eastern states) who identify to varying degrees as being a conservationist.

### 5. Attitudes about cougar restoration and political identity

Survey participants were asked to indicate their political orientation on a 7-point scale ranging from very liberal to moderate to very conservative. In the graph below (Figure 7), the labeled line shows the association between one’s political orientation and attitudes toward cougar restoration.

**Figure 7.**
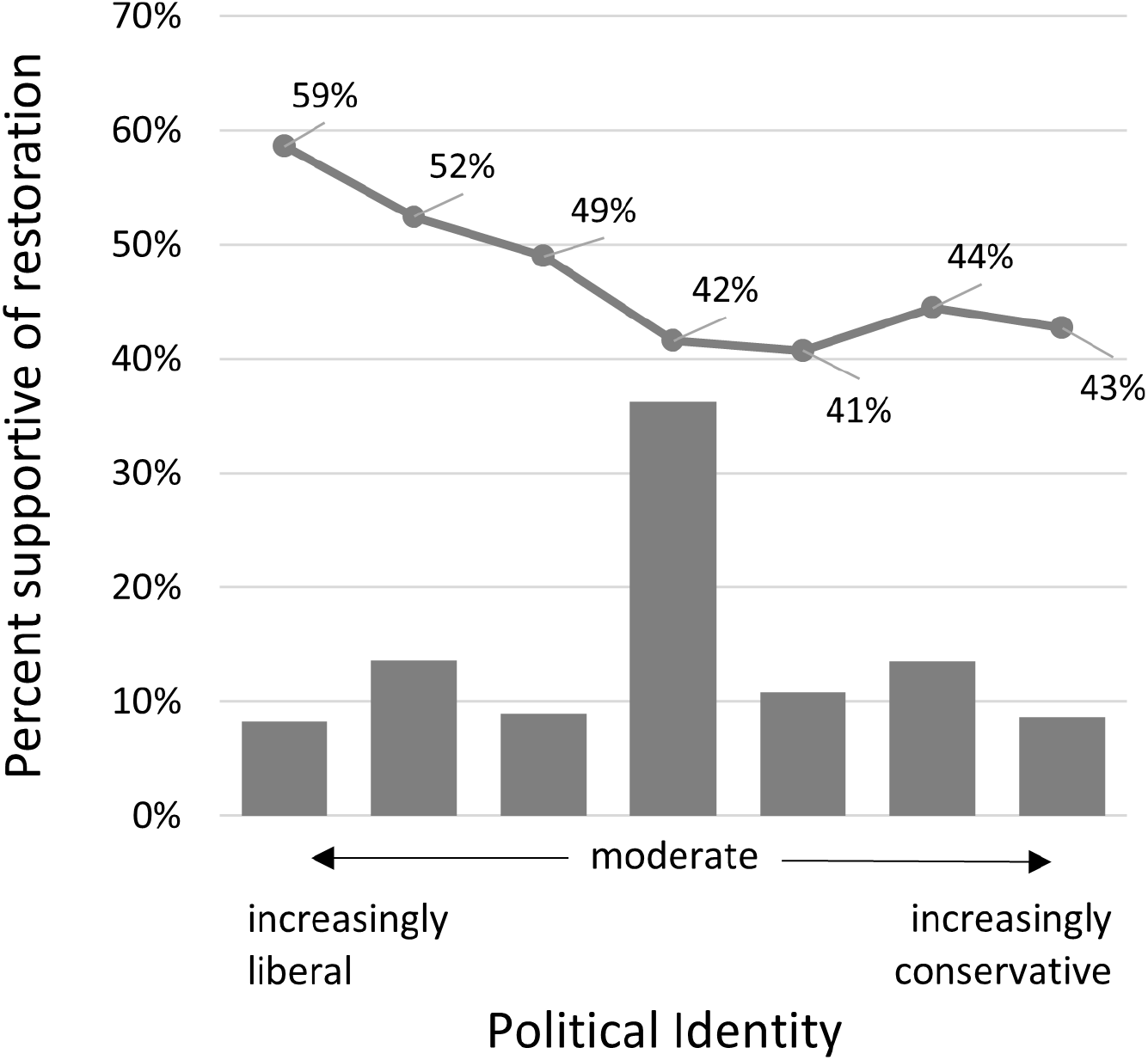
Attitudes toward cougar restoration by political identity.

The data indicate that attitudes about cougar restoration are similar among moderates and conservatives (about 42% supportive). Furthermore, support for cougar restoration increases as one’s political orientation becomes increasingly liberal, with the most liberal people being nearly 60% supportive of cougar restoration.

The lower panel of this graph provides additional context by indicating the portion of the sample (for the seven eastern states) who identify to varying degrees as being liberal, moderate, or conservative.

### 6. Attitudes about cougar restoration and living environment

Survey participants were asked to indicate the size of the community where they live. The list of responses from which a survey participant could select was:

- Large city with 250,000 or more people
- City with 100,000 to 249,999 people
- City with 50,000 to 99,999 people
- Town with 10,000 to 24,999 people
- Town with 5,000 to 9,999 people
- Small town/village with less than 5,000 people
- A farm or rural area

The graph to the right (Figure 8A) suggests that attitudes toward cougar restoration vary slightly among the different-sized communities. The lower panel of this graph (Figure 8B) provides additional context by indicating the portion of the sample (for the seven eastern states) who live in different-sized communities.

**Figure 8.**
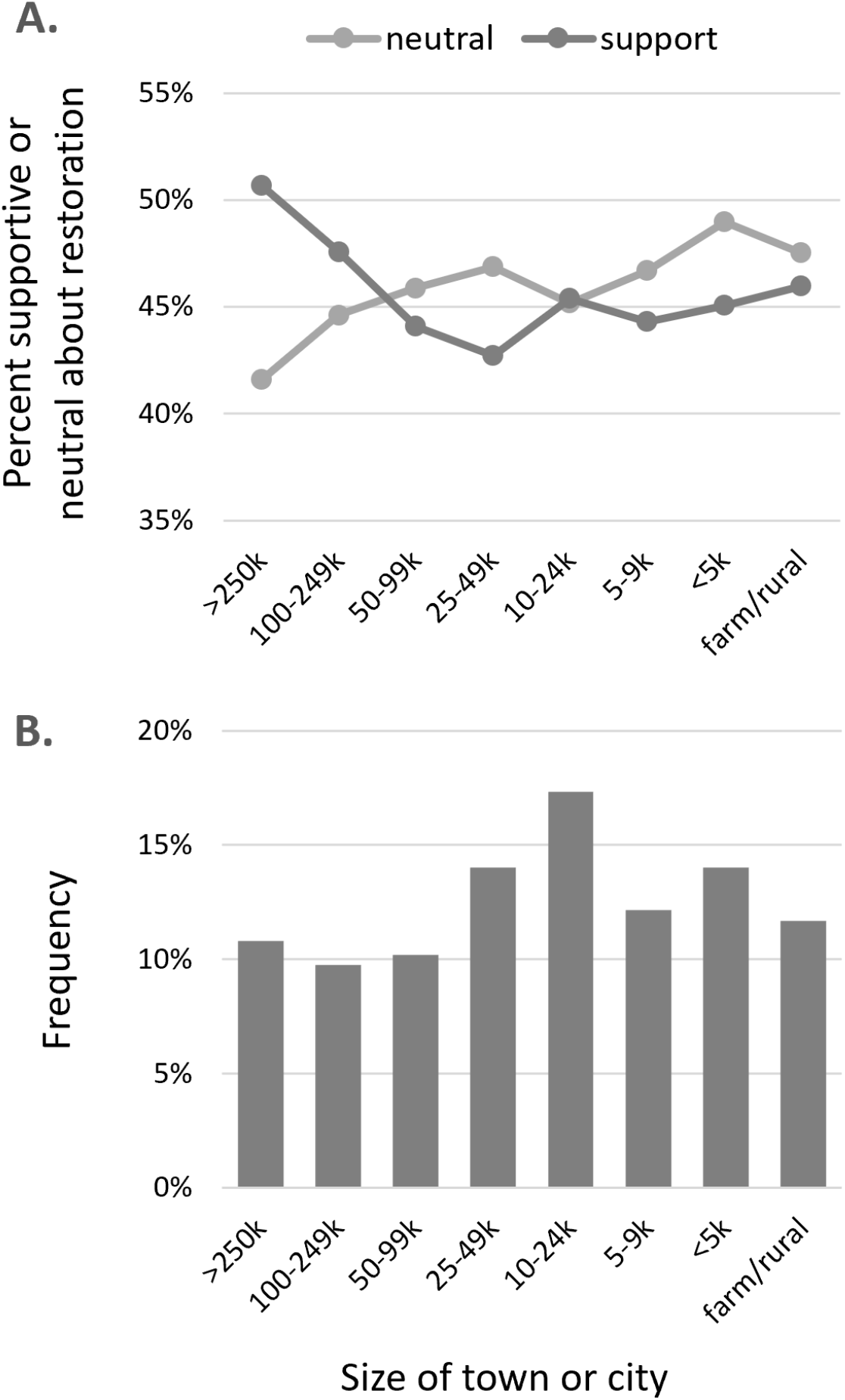
Support for cougar restoration by residency size.

### 7. Attitudes about cougar restoration and wildlife value orientations

Survey participants were asked to respond to a set of items designed to collectively represent their beliefs about how humans should relate with wildlife. These items are designed to capture two types of value-orientations, *mutualism* and *domination*. Domination represents the idea that wildlife are subordinate to humans and should be used in ways that benefit humans; mutualism represents the idea that wildlife are morally relevant parts of one’s social community, deserving of care and compassion.

Nine items are used to measure mutualism, and 10 items are used to measure domination. Each item has a response set that is a 7-point bipolar scale (agree/disagree). We calculated a score for each participant as the average response to each set of items. Then we binned responses according to this conversion table:

| Actual Score | [1, 1.5] | (1.5, 2.5] | (2.5, 3.5] | (3.5, 4.5] | (4.5, 5.5] | (5.5, 6.5] | (6.5, 7] |
| --- | --- | --- | --- | --- | --- | --- | --- |
| Binned score | 1 | 2 | 3 | 4 | 5 | 6 | 7 |

Figures 9A and 9B show how one’s mutualism score (Figure 9A) and domination score (Figure 9B) are associated with their attitude about cougar restoration. The red circles are mean responses on a scale of [-2,2], where 0 is neutrality, positive values are support for restoration, and negative values are opposition. The bars mark the 95% confidence intervals (CIs; calculated as two times the standard error).

**Figure 9.**
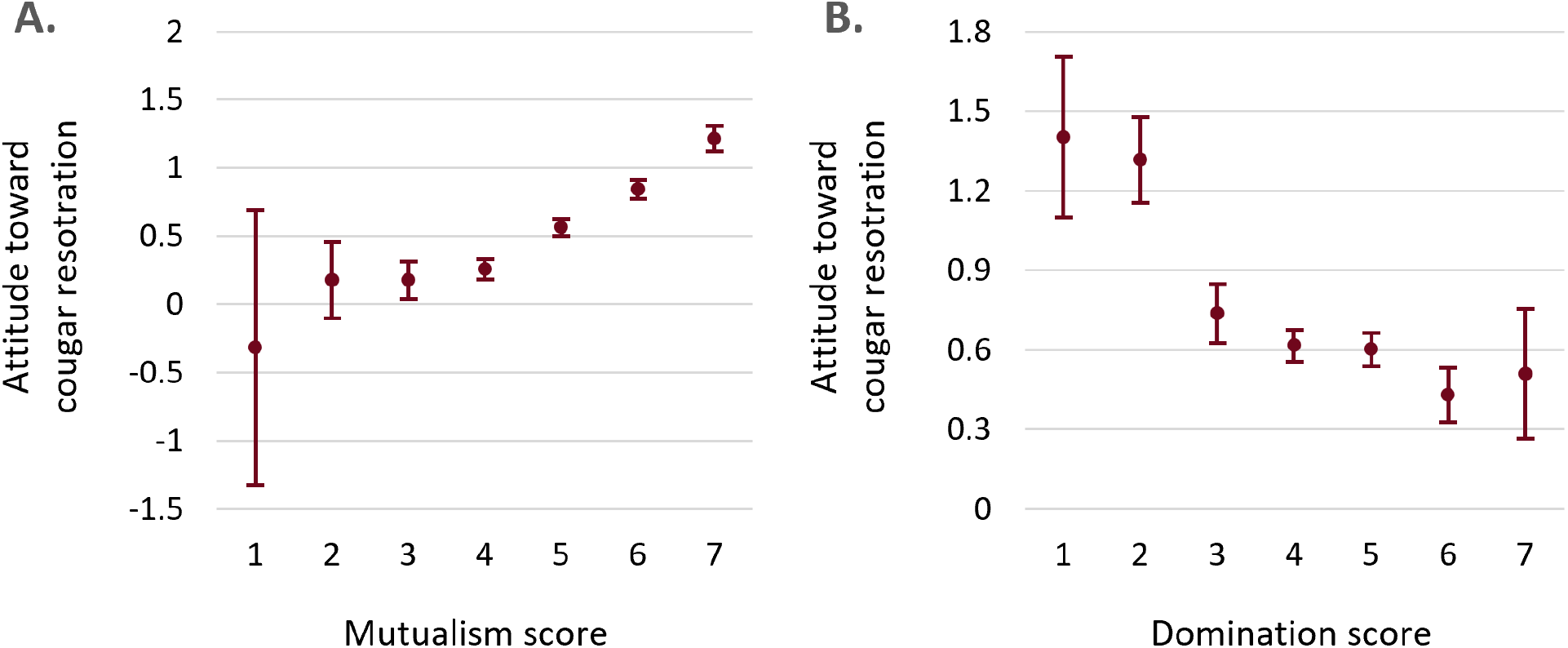
Attitudes toward cougar restoration in relationship to one’s scores for the mutualism (left) and domination (right) value orientations.

Figures 10A and 10B show the frequency distribution of mutualism and domination scores in the sample. For example, the most common binned score for mutualism was 5, which was nearly 30% of the sample.

**Figure 10.**
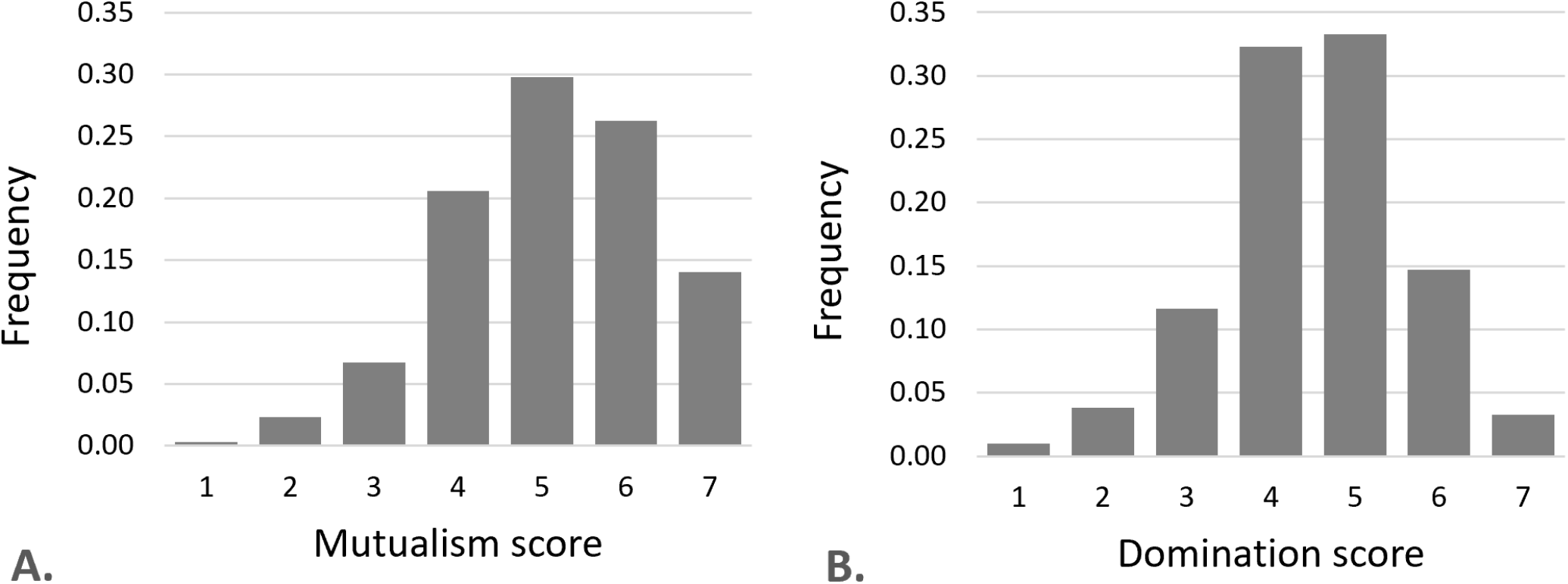
Distribution of scores measuring mutualism and domination.

For context: that mutualism and domination scores are so highly predictive of attitudes toward cougar restoration is importantly a statistical consequence of having binned those scores. One can also perform multiple linear regression (n=2756) where the predictors are the actual scores. In that case, the actual scores explain approximately 12% of the variance in attitudes toward restoration.

### 8. Attitudes about cougar restoration and gender

The graph to the right (Figure 11) shows that males are, on average, more supportive of cougar restoration than females. In that graph, the circles are means and the vertical bars mark 95% CIs (calculated as two times the standard error).

While we did not expect there to be a difference between males and females, a difference is not surprising. The most likely explanations are: (i) the idea of carnivores is more attractive to men, (ii) women perceive greater risks associated with carnivores, or (iii) some combination of those two phenomena.

**Figure 11.**
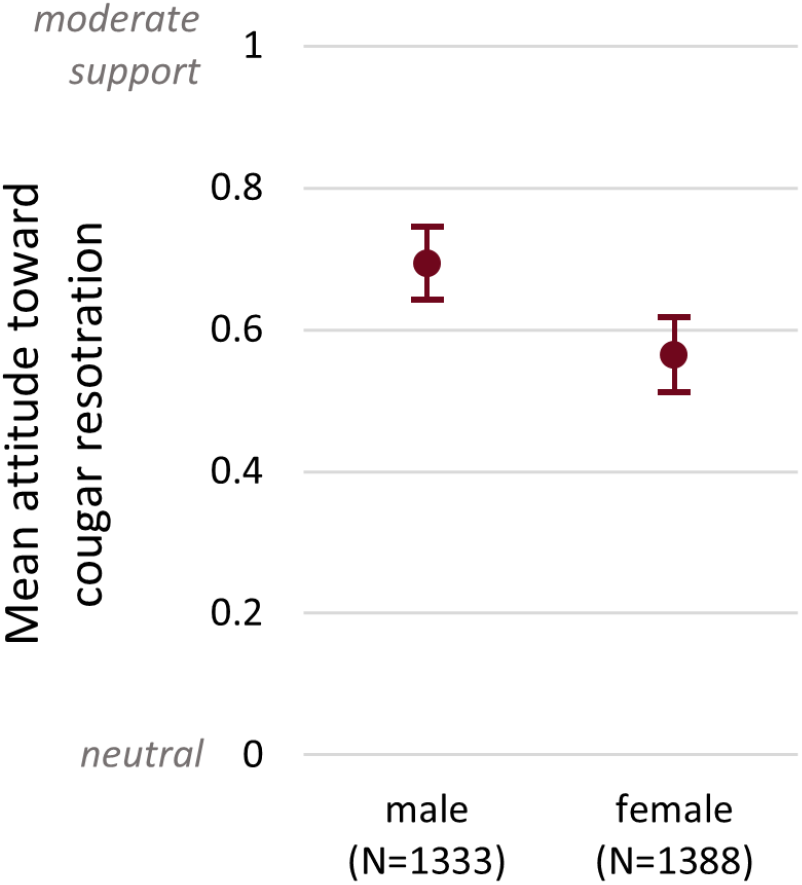
Attitudes toward cougar restoration by gender.

### 9. Attitudes toward cougar restoration and behavioral intentions

Participants were presented with a list of actions they could take in support of or opposition to cougar restoration in their state of residence. Participants were asked to indicate how likely or unlikely they were to engage in each action (i.e., their behavioral intentions). The complete list of actions included:

- Contact a wildlife manager/management agency
- Write your Congressperson
- Post to Facebook or Twitter
- Contribute to an organization
- Sign a petition
- Shoot a cougar if you saw one *(included as an oppositional action only)*

Approximately half of respondents reported a likelihood of engaging in at least one action in support of cougar restoration. Figure 12 shows the frequency of respondents who reported they were “likely” or “very likely” to engage in each behavior.

**Figure 12.**
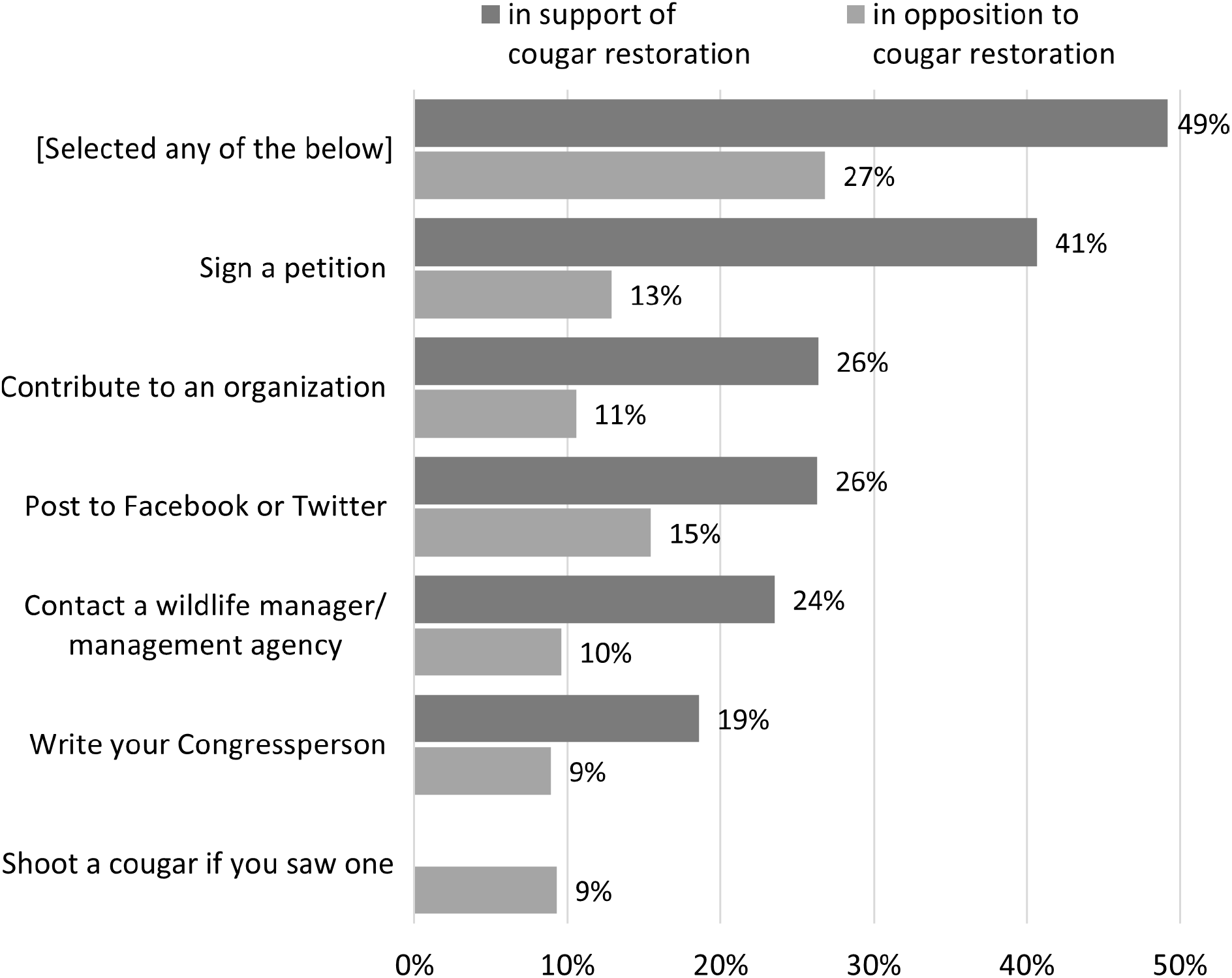
Intention to engage in actions supporting or opposing cougar restoration.

### 10. Attitudes toward cougar restoration on a state-by-state basis

The state-specific attitudes toward cougar restoration are shown below in Figure 13. Results in this section are based on responses to a single survey
item: “If cougars were reintroduced to rural areas of the eastern United States, what would be your reaction?”. We chose to present this survey item because it exhibited the greatest proportion of respondents who reported strong attitudes (support or opposition) to cougar restoration, so provides the best gauge of participants’ leanings when it comes to the issue of cougar restoration. We provide the same state-level results for the other survey item and a combination of the two items in <u>Appendix A</u>.

**Figure 13.**
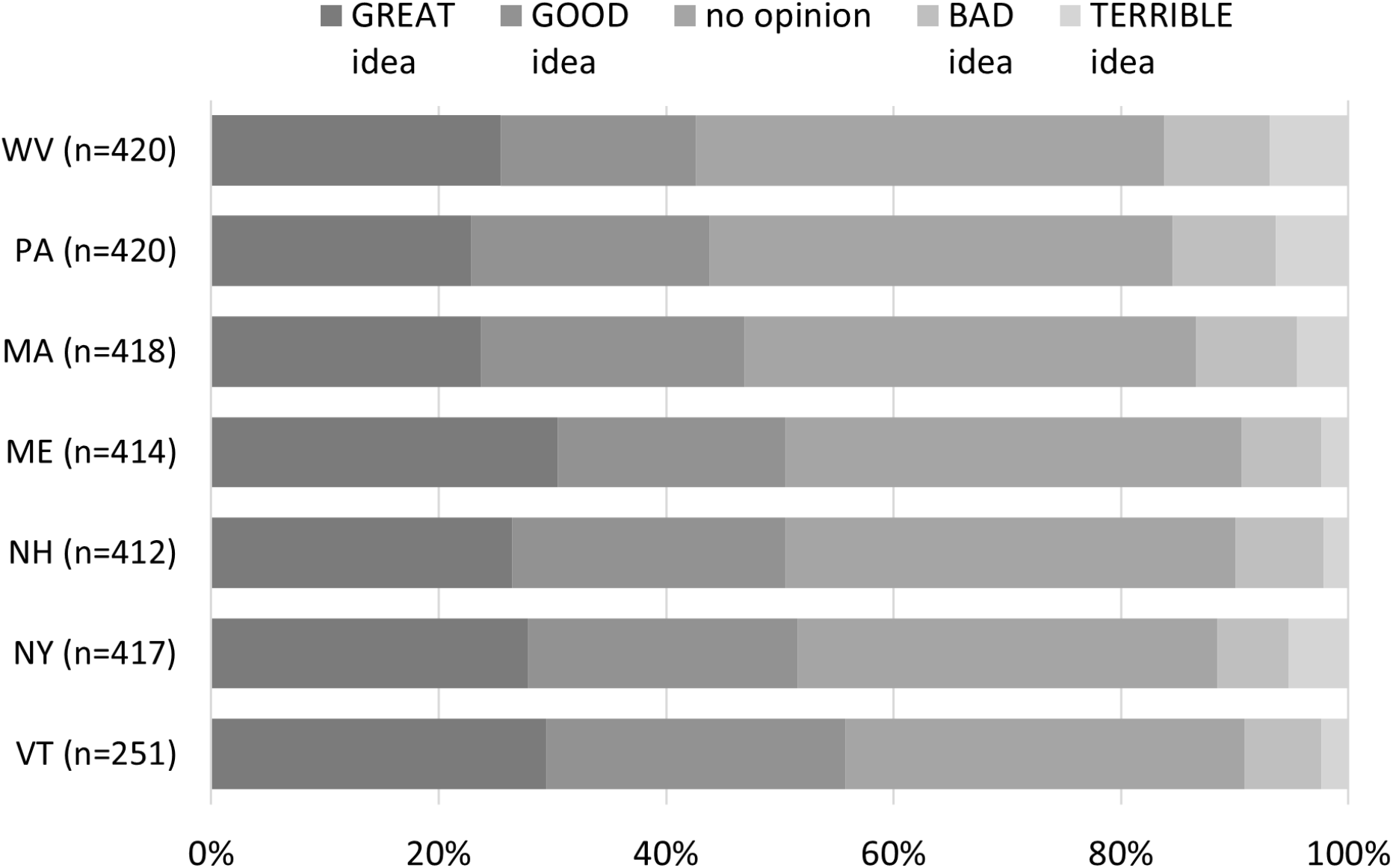
Attitudes toward cougar restoration by state.

While there are state-level differences in attitude, the differences are not readily apparent with this presentation of the data. To better facilitate comparisons among states, we conducted additional analyses. In particular, Figure 14 (next page) shows the percentage of participants expressing strong support (horizontal axis) in relation to the ratio of strong support to strong opposition (vertical axis). To better understand the meaning of that ratio, consider New York which has a ratio of 5. This can be interpreted to mean, for every person who might strongly oppose cougar restoration one might expect approximately five people to strongly support cougar restoration.

**Figure 14.**
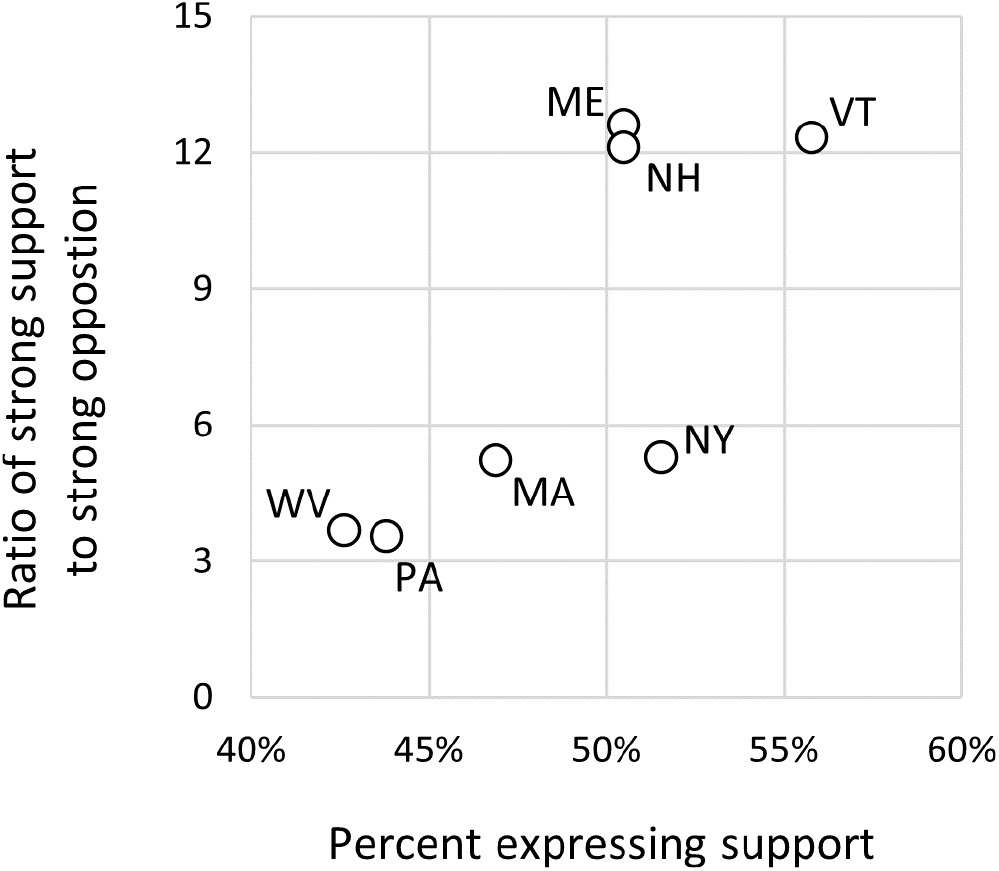
Ratio of strong support to strong opposition in relationship to percent expressing support.

Generally speaking, the intensity of support for cougar restoration increases as one moves to upper, right portions of Figure 14.

The horizontal axis of the graph to the below (Figure 15) is the same as the previous graph. But here, the vertical axis represents the proportion of the sample expressing “no opinion” about cougar restoration. The relevance of the vertical axis is that people expressing “no opinion” would generally be thought to be more likely (than people expressing support or opposition) to develop a supportive attitude if given positive information about cougar restoration and develop an oppositional attitude if given negative information.

**Figure 15.**
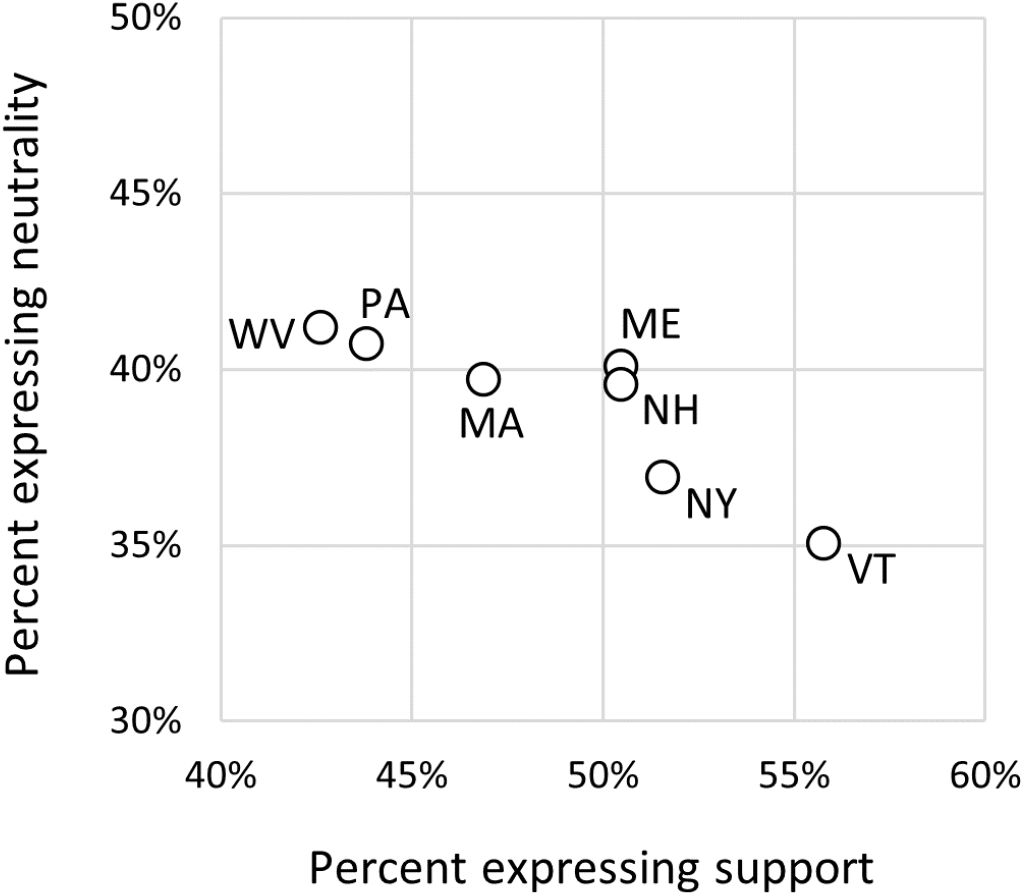
Percent expressing neutrality in relationship to percent expressing support.

As one moves from lower to upper portions of Figure 15 there is an increasing portion of a state’s population whose view is most likely to change if given additional information about cougar restoration.

Note about figure 14 and 15: Strong support refers to people who thought restoration was a “GREAT idea.” Strong opposition refers to people who thought restoration was a “TERRIBLE idea.” Support refers to people who thought cougar restoration was a “GREAT idea” or a “GOOD idea.”

Figure 16 (to the left) uses a map to depict the same state-level spatial variation in attitudes toward cougar restoration that is presented in Figure 14. The ratios (bold font) represent the ratio of strong support to strong opposition, and the percentages (italicized font) represent the percentage of respondents indicating strong support for cougar restoration.

**Figure 16.**
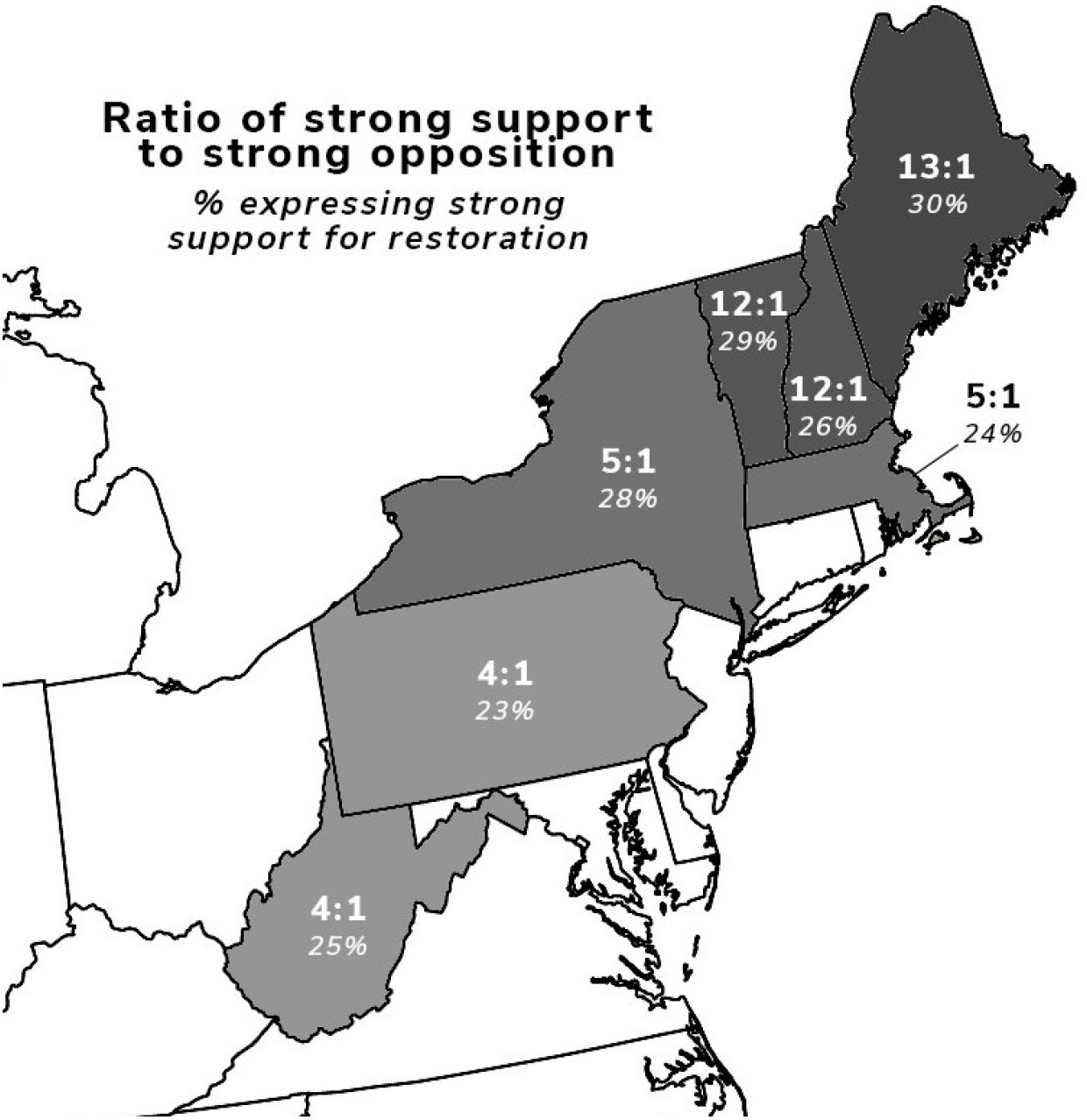
Map of two metrics of support for cougar restoration.

## CONCLUSION

The primary objective of this study was to provide a preliminary assessment of attitudes toward cougar restoration for seven eastern U.S. states with potential cougar habitat (ME, MA, NH, NY, PA, VT, and WV). Results from an online survey of residents within these states (n=2756) suggest that support for cougar restoration is much higher than opposition to cougar restoration. Ratios of strong support to strong opposition range from approximately 4:1 to 13:1. Maine, Vermont, and New Hampshire have the highest ratio of strong support to strong opposition (13:1, 12:1, and 12:1 respectively) while Pennsylvania and West Virginia have the lowest ratio of strong support to strong opposition (4:1 each). Results may prove especially useful for identifying and prioritizing potential sites for the restoration of cougars within their historic eastern range.

## ACKNOWLEDGEMENTS

This study was funded for the Cougar Research Collaborative (CRC) by the Northeast Wilderness Trust, Panthera and Tompkins Conservation. We also wish to thank members of the CRC who provided invaluable feedback on this study. Specifically, we wish to thank Mark Elbroch, Shelby Perry, Tom Butler, Lydia Roe, and Michael Manfredo for their helpful feedback.

## APPENDIX A

**Table A1.**
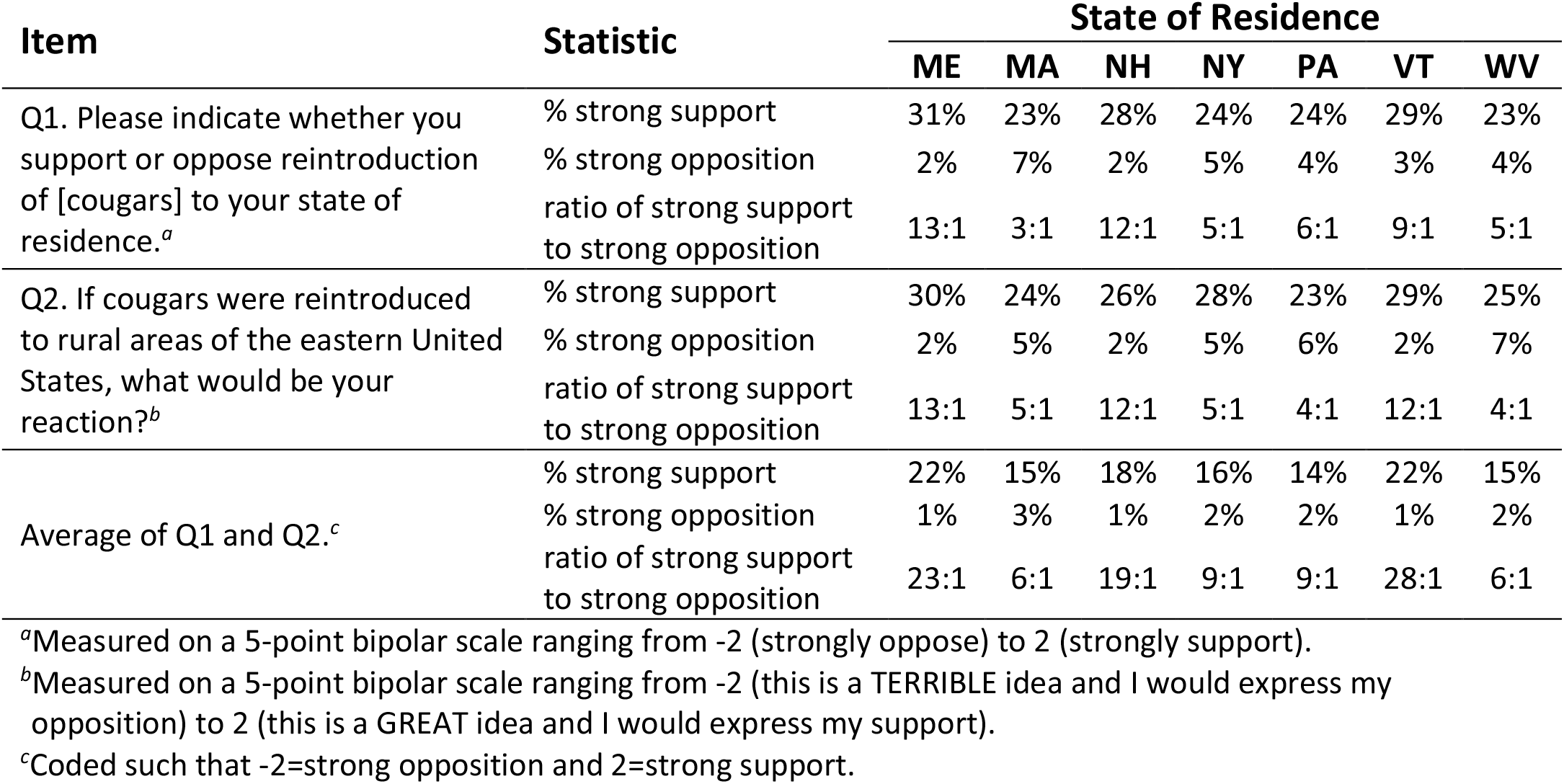
State-level summary of items used to measure attitudes toward cougar restoration.

| Item | Statistic | State of Residence |  |  |  |  |  |  |
| --- | --- | --- | --- | --- | --- | --- | --- | --- |
|  |  | ME | MA | NH | NY | PA | VT | WV |
| Q1. Please indicate whether you support or oppose reintroduction of [cougars] to your state of residence.<sup>a</sup> | % strong support | 31% | 23% | 28% | 24% | 24% | 29% | 23% |
|  | % strong opposition | 2% | 7% | 2% | 5% | 4% | 3% | 4% |
|  | ratio of strong support to strong opposition | 13:1 | 3:1 | 12:1 | 5:1 | 6:1 | 9:1 | 5:1 |
| Q2. If cougars were reintroduced to rural areas of the eastern United States, what would be your reaction?<sup>b</sup> | % strong support | 30% | 24% | 26% | 28% | 23% | 29% | 25% |
|  | % strong opposition | 2% | 5% | 2% | 5% | 6% | 2% | 7% |
|  | ratio of strong support to strong opposition | 13:1 | 5:1 | 12:1 | 5:1 | 4:1 | 12:1 | 4:1 |
| Average of Q1 and Q2.<sup>c</sup> | % strong support | 22% | 15% | 18% | 16% | 14% | 22% | 15% |
|  | % strong opposition | 1% | 3% | 1% | 2% | 2% | 1% | 2% |
|  | ratio of strong support to strong opposition | 23:1 | 6:1 | 19:1 | 9:1 | 9:1 | 28:1 | 6:1 |

<sup>a</sup>Measured on a 5-point bipolar scale ranging from -2 (strongly oppose) to 2 (strongly support).
<sup>b</sup>Measured on a 5-point bipolar scale ranging from -2 (this is a TERRIBLE idea and I would express my opposition) to 2 (this is a GREAT idea and I would express my support).
<sup>c</sup>Coded such that -2=strong opposition and 2=strong support.

